# The infant gut virome assembles rapidly and predictably over the first three years of life

**DOI:** 10.64898/2026.02.17.705938

**Authors:** Michael Shamash, Corinne F. Maurice

## Abstract

Assembly of the gut microbiota in infancy is an important health determinant, yet the viral component, the virome, remains poorly described. Here, we conduct a meta-analysis of 12 infant cohorts across 8 countries to model gut virome development over the first three years of life. Our results revealed distinct diversity patterns, where viral richness increased significantly over the first eight months of life alongside a loss of community evenness. We define virome developmental velocity, the rate of change between sequential samples, which significantly decreased over time. This was mirrored by community-wide convergence, as infant viromes became more similar with age. Functionally, the transition to a stable state involved a significant decrease in extracellular temperate phages and shifts in phage-encoded auxiliary metabolic genes, which can modulate host metabolism. Our work provides a baseline characterization of gut virome development, marked by a conserved successional trajectory, providing a framework to identify disease-associated perturbations.

## Introduction

The human gut microbiota begins its development at birth which continues throughout the first few years of life (1, 2). Many studies have focused on the gut bacteriome and its associations with infant health (3–6). While several studies have begun to shine light on the early life gut virome, primarily composed of bacteria-infecting viruses known as bacteriophages (phages), much remains unknown about these highly abundant members of the gut microbial ecosystem (7–10). Current evidence suggests that this viral community is not randomly acquired but instead linked to the arrival of the first bacterial hosts. The initial phages in the gut are induced prophages from pioneer gut bacteria and remain as prominent members of the gut virome for several months (11). While bacterial colonizers follow a predictable successional trajectory (12), it remains unclear whether assembly of the gut virome follows a similar structured assembly.

The gut nutritional landscape undergoes important shifts during infancy, primarily driven by the transition from a diet rich in human milk oligosaccharides (HMOs) to one which is complex and contains plant-derived polysaccharides (13). While the gut bacteriome adapts to these shifts through changes to its structure and composition, the role of the virome in supporting this transition remains unexplored. Indeed, phages can carry auxiliary metabolic genes (AMGs) which modulate host metabolism (14). These genes are best characterized in aquatic and soil ecosystems, where phages can carry photosynthetic genes (15); however, studies have also identified enrichment in nitrogen and carbon utilization genes among phages in the adult gut (16–18). AMGs found in the gut virome during early life have yet to be characterized, leaving a gap in our understanding of how phages may actively contribute to shifts in bacterial metabolism during this important developmental window.

Defining the healthy developmental program of the infant gut virome is an important step in identifying alterations linked with disease; however, a major hurdle is the extreme sequence-level variation between individuals, where two individuals may share very few viral sequences (19). To address this, we recently proposed a novel analytical approach to allow for improved inter-study and inter-individual virome comparisons (20). By grouping viral sequences based on their predicted bacterial host family (PHF), we can overcome the highly individualized nature of gut viromes to identify broader associations based on phage functional potential.

Using this approach, we conduct a meta-analysis of infant gut viromes to identify global patterns in phage community assembly and functional capacity. We focus on published studies which have free virome (or virus-like particle-enriched) sequencing data, rather than bulk metagenomes. This approach allows for the detection of rarer viruses often missed in complex bulk metagenomes (21), despite reducing the number of eligible studies. This focus on the extracellular, actively replicating virome captures the viral fraction most likely driving microbial community assembly dynamics. We identify 12 studies from 8 countries for analysis, consisting of 1,893 samples from 999 infants (11, 22–32). Most of these studies are limited by small sample sizes or focus on specific geographic regions, while also using distinct analytical approaches. By employing a unified bioinformatic pipeline across these 12 diverse cohorts, we mitigate technical biases to reveal a conserved, global developmental program of the infant gut virome. We describe the development of the gut virome from birth to three years of age, finding that despite rapid fluctuations in the first few months of life, the extracellular virome converges to a stable state by two years of age. This transition from a highly dynamic to stable community is also characterized by a decrease in the abundance of extracellular temperate phages. Using a regression model, we accurately predict infant age from gut virome composition, identifying associations between groups of phages and infant chronological age. Finally, we observe enrichment in certain metabolic pathways among certain groups of phages, suggesting that the virome’s functional potential changes in response to major developmental milestones.

## Methods

### Study curation

The following inclusion criteria were used to identify studies to be included in our meta-analysis: (1) public availability of Illumina sequencing data derived from the fecal extracellular virome (enriched virus-like particles); (2) sampling within the first three years of life (0-36 months); and (3) a “healthy” clinical status as defined by the original study investigators. For datasets containing both healthy and clinical cohorts (e.g. type I diabetes or inflammatory bowel disease), only the healthy controls were retained for analysis. This selection process yielded 12 distinct studies, comprising six longitudinal (11, 22–24, 28, 29) and six cross-sectional (25–27, 30–32) studies (**Table S1**).

### Raw data processing and viral OTU (vOTU) library construction

Raw sequencing reads retrieved for each study from the NCBI Sequence Read Archive (SRA) or European Nucleotide Archive (ENA) using study-specific accession numbers (**Table S1**). Read quality control was performed using fastp (v0.23.4) (33) with the following parameters: -q 15 --cut_right --cut_window_size 4 --cut_mean_quality 20 --length_required 50. For paired-end datasets, the --detect_adapter_for_pe parameter was added. To remove host contamination, reads were mapped against the *Homo sapiens* GRCh38 reference using bowtie2 (v2.5.2) (34) in sensitive mode.

Cleaned reads were assembled *de novo* for each sample individually using metaSPAdes (v3.15.4) (35) with default parameters. Viral sequences were identified from the resulting contigs using geNomad (v1.7.4) (36) in end-to-end mode. To generate a non-redundant viral catalogue, viral contigs were dereplicated into viral operational taxonomic units (vOTUs). An all-versus-all BLASTN (v2.14.0) (37) search was performed, and contigs were clustered using anicalc.py and aniclust.py scripts from CheckV (38) based on ≥95% average nucleotide identity (ANI) over 85% of the shorter contig’s length. vOTUs shorter than 3kb were excluded from the final library. To quantify vOTU abundance, filtered reads from each sample were mapped back to the vOTU library using Bowtie2 (v2.5.2) (34). Coverage statistics were generated using samtools (v1.18) (39). A vOTU was considered “detected” in a sample only if it met a minimum threshold of ≥1X mean depth of coverage and ≥75% breadth of coverage (40). Computational host prediction was performed using iPHoP (v1.0.0, db version: Sept_2021_pub) (41), and viral lifestyles (temperate or non-temperate) were predicted using BACPHLIP (v0.9.6) (42).

### Alpha diversity analyses and statistical modelling

Alpha diversity metrics (observed richness, Shannon index, and Pielou’s evenness) at the vOTU level were calculated in R (v4.5.1) using phyloseq (1.52.0) (43). To model diversity trajectories while controlling for inter-study heterogeneity and repeated measures, generalized additive mixed models (GAMMs) were implemented with mgcv (v1.9-3) (44). Different model distributions were chosen based on the properties of each diversity metric. Observed richness was modelled using a negative binomial distribution (family = nb()) to account for overdispersed count data. Shannon index was modelled using a Gaussian distribution (family = gaussian()). Pielou’s evenness was modelled using a beta regression (family = betar()) to accommodate continuous values constrained in the (0, 1) interval. For Pielou’s evenness, values were transformed prior to modelling to compress the [0, 1] interval to (0, 1) following the formula 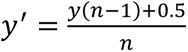, as described by Smithson and Verkuilen (45), where n is the total number of samples, to accommodate the requirements of the beta distribution.

The general model formula was defined as y ∼ s(age_months) + log10(depth) + s(age_months, study, bs = “fs”, m = 1) + s(subjectID, bs = “re”). Library size (log_10_ transformed) was included as a fixed effect to control for varying sequencing depth across samples without the need for rarefaction. To maximize statistical power and maintain the maximum possible sample size, other clinical metadata variables (infant sex and delivery mode) were not considered in the models, as complete information for these variables was only available for 401 of the 1,893 total samples (21%).

Overall model performance was evaluated based on deviance explained and the significance of the smooth terms. Visualization of the fitted smooth terms was performed using the gratia package (v0.11.1) (46).

To identify the specific window where observed richness was increasing before stabilizing, we analyzed the first derivative of the fitted age_months smooth term using the derivatives() function from the gratia package. The point of stabilization was defined as the earliest age (in months) where the 95% confidence interval of the first derivative became negative, indicating the rate of change of richness was no longer significantly different from zero.

### Prevalence of vOTUs and PHFs

To assess the distribution of viruses across cohorts, we calculated the prevalence of both individual vOTUs and phage host families (PHFs). PHFs are groupings of vOTUs based on their predicted bacterial host family. We have previously shown that PHFs are a functionally relevant, qualitative unit of phage classification which can improve virome comparisons across studies (20).

Samples were grouped into five age bins: 0 months (neonatal), 1-3 months (early infancy), 4-6 months (transitional), 7-12 months (late infancy), >1 year (toddler). These intervals were selected based on both sampling density and their alignment with key infant developmental milestones, specifically the transition from exclusive milk or formula feeding (0-3 months) through the introduction of solid foods (4-6 months) to being fully weaned on a diversified diet (>12 months). Prevalence was defined as the proportion of infants within an age bin which have a vOTU/PHF at detectable levels (see above). For longitudinal studies where an infant was sampled multiple times within an age bin, the vOTU/PHF was considered present if detected in at least one sample during that period. This binning strategy was used to standardize the contribution of infants from both cross-sectional and longitudinal cohorts.

### Beta diversity analyses

Beta diversity was assessed using the weighted UniFrac metric on PHF-level viromes. The GTDB tree used by iPHoP (bac120_r202.tree) was used for weighted UniFrac distance calculations. To test the impact of metadata variables (e.g., infant age, study) on viral community composition, a permutational multivariate analysis of variance (PERMANOVA) test was performed using the adonis2() function in vegan (2.7-2) (47). The primary model included library size (log_10_ transformed), study, and age (months). To account for the longitudinal nature of the data, permutations were constrained within individual subjects using the strata argument. A second PERMANOVA model was used to quantify the variance explained (R^2^) by inter-individual differences. This model’s formula included only library size (log_10_ transformed), study, and subjectID. No p-value is reported for the subjectID term as the high number of levels (n = 991) relative to the total number of samples prevents us from conducting a valid permutation test for this variable, rather it is included only to estimate the total R^2^ of subjectID.

### Virome developmental velocity and beta dispersion

To quantify the rate of virome community change over time, we calculated virome developmental velocity using the six longitudinal studies (n = 989 samples from 120 individuals, across 6 studies). Developmental velocity was defined as the weighted UniFrac distance between consecutive samples from the same infant, divided by the time interval (in months) between these consecutive samples. The age assigned to each velocity estimate was that of the second (later) sample. Inter-individual variation was assessed with beta dispersion, calculated as the distance of each sample to its age-specific (in months) centroid in multivariate space, using the betadisper() function in the vegan package.

Both velocity and beta dispersion metrics were modelled using GAMMs with a Tweedie distribution (family = tw()) to accommodate the non-negative nature of the data. The velocity model included a fixed effect for mean_log10_depth (the average library size of the two compared samples), while the beta dispersion model included log10(depth). Both models used the previously described fixed and random effect structure to account for heterogeneity among cohorts: y ∼ s(age_months) + mean_log10_depth + s(age_months, study, bs = “fs”, m = 1) + s(subjectID, bs = “re”). To maximize statistical power and maintain the maximum possible sample size, other clinical metadata variables (infant sex and delivery mode) were not considered in the models, as complete information for these variables was only available for 401 of the 1,893 total samples (21%). Visualization of the fitted smooth terms was performed using the gratia package.

### Random forest age predictor

To determine the predictability of virome maturation, we trained a random forest (RF) regression model using the randomForest (v4.7-1.2) (48) package to predict infant age (months) based on PHF relative abundances. The dataset was split into training (70%) and testing (30%) sets using the createDataPartition() function from the caret (v7.0-1) (49) package to maintain a balance of age across both datasets. Model performance was evaluated by calculating the root mean squared error (RMSE) and variance explained (R^2^) on both training and test datasets. To evaluate the model’s accuracy over the course of infant development, a sliding window approach was used, where RMSE was calculated for overlapping 3-month age windows. This higher-resolution approach provides a continuous assessment of the model’s predictive performance during periods of rapid virome change, avoiding boundary effects which could arise when using the discrete age bins used for prevalence calculations.

The most important features (PHFs) were identified using the IncNodePurity metric from the importance() function, part of the randomForest package. A heatmap of the top 20 most important PHFs was generated using ComplexHeatmap (v2.24.1) (50) to visualize their mean abundance patterns during the first years of life. For each PHF, a Pearson correlation coefficient (r) was calculated against age_months. These values were squared to determine the coefficient of determination (R^2^), which represents the proportion of variance in age explained by the individual PHF. A signed R^2^ value was calculated by multiplying the R^2^ by the sign of the correlation coefficient, to indicate the direction of the PHF’s association with age.

### Auxiliary metabolic gene identification and analysis

To characterize the functional potential of the infant virome, protein-coding sequences were predicted from the vOTU library using prodigal-gv (v2.11.0) (36, 51) in meta mode. The resulting gene set was clustered into a non-redundant catalog using mmseqs2 (b804fbe384e6f6c9fe96322ec0e92d48bccd0a42) (52) based on ≥95% average amino acid identity (AAI) and ≥ 90% coverage.

Functional annotation was performed by searching the non-redundant gene catalog against the KOfam database (v20250201) using hmmsearch (HMMER v3.4) (53) with an E-value threshold of 1 × 10^−5^. Results were processed in R using the rhmmer package (v0.2.0) (https://github.com/arendsee/rhmmer). Auxiliary metabolic genes (AMGs) were specifically identified by filtering annotations against a curated list of KEGG orthologs (KOs) established by the virus identification tool VIBRANT (54).

A heatmap of AMG metabolism categories was generated using pheatmap to visualize abundance patterns during the first years of life. AMG abundance (either at the metabolism or pathway level) within each sample was calculated as the cumulative abundance of all vOTUs carrying that specific AMG. To avoid biases introduced by potential gene duplications or functional redundancy, copy number was not considered. Relative abundances were min/max-scaled per category to facilitate comparisons of trends across categories with significantly differing abundances.

Enrichment was assessed using a hypergeometric test (Fisher’s exact test). This test compares the frequency of a specific AMG category within a single PHF against its global frequency across all PHFs. By accounting for the number of unique vOTU within each PHF, this test determines if a functional category is significantly overrepresented relative to the size of that PHF. P-values were adjusted for multiple testing using the Benjamini-Hochberg method, and categories were considered significantly enriched if the adjusted p-value was less than 0.05.

## Results

### Integrating 12 cohorts to study the healthy infant gut virome

To characterize the developmental dynamics of the gut virome during early life, we curated a dataset composed of 12 independent studies across eight countries (**Figure 1A, Table S1**). All included studies used viral particle enrichment protocols, such as filtration or ultracentrifugation, to sequence the extracellular virome rather than bulk metagenomes, ensuring a high-resolution view of the viral community. The combined dataset included 1,893 enriched fecal viromes from 999 healthy infants, from birth (during the first few days of life) to 38 months of age (**Figure 1B**). All raw sequencing data were re-processed through a standardized bioinformatic pipeline (**Figure 1C**) to ensure cross-study comparability. This involved *de novo* assembly of individuals samples, viral identification, and generation of a non-redundant viral operational taxonomic unit (vOTU) catalog containing 49,745 sequences ≥ 3kb in length. Following read mapping and the application of coverage detection thresholds (≥1X depth and ≥75% breadth of coverage), 1,869 samples from 991 infants were retained for downstream analysis (24 samples from 8 infants had insufficient sequencing depth for analysis). While sequencing depth varied across studies (mean 10,494,039 ± 27,272,299 raw reads [1.165 ± 2.63 Gbp] per sample), library size was accounted for in all downstream modelling to control for potential sampling bias.

**Figure 1.**
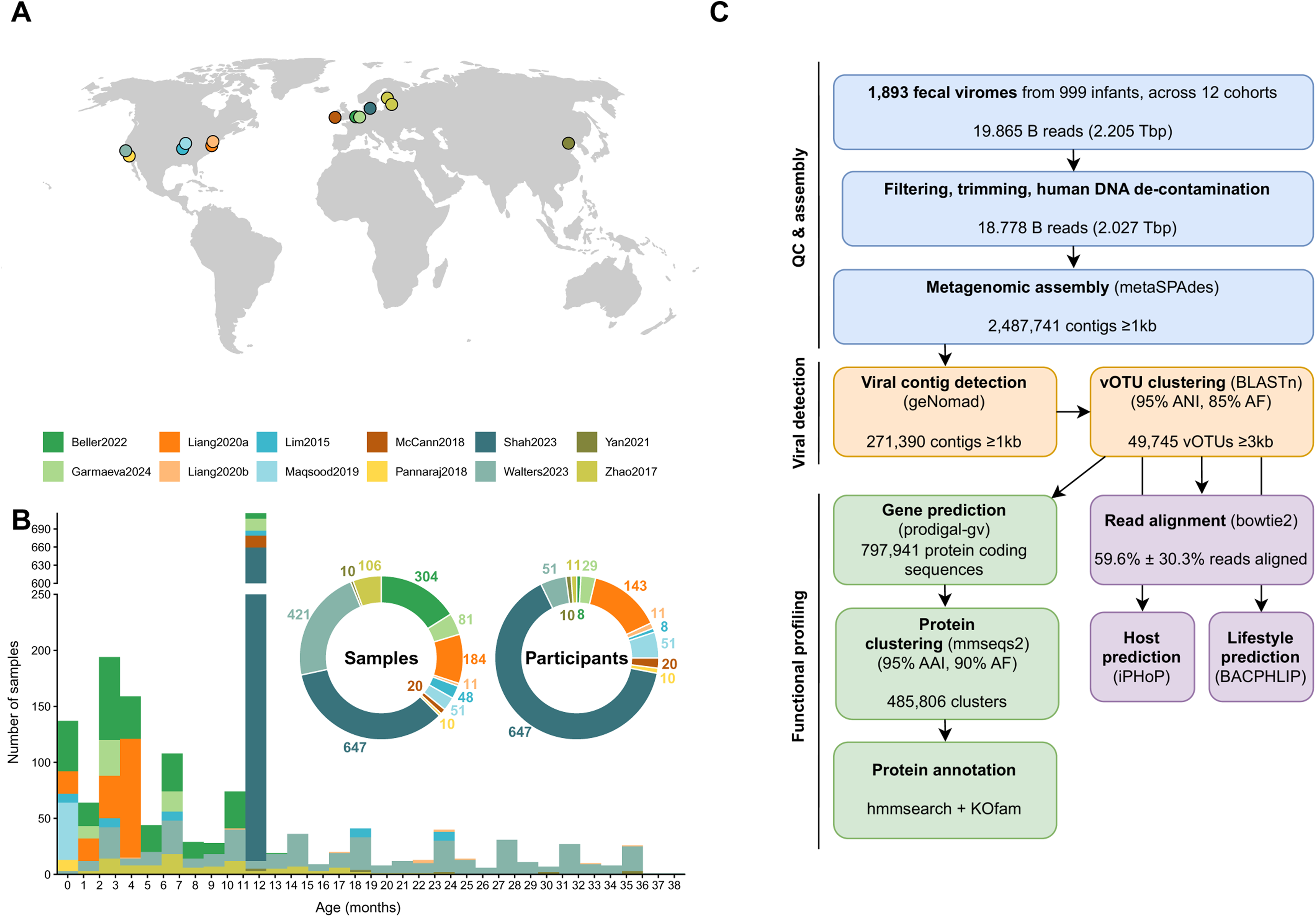
Integrated analysis and characterization of the global infant gut virome. **(A)** Geographic overview of sampling locations for the 12 cohorts included in this meta-analysis, representing 999 infants (11, 22–32). **(B)** Longitudinal sampling distribution across the first three years of life. Inset plots summarize the total number of samples (n = 1,893) and participants contributed by each study (n = 999). **(C)** Schematic of the bioinformatic pipeline. A total of 19.865 billion reads were processed through quality control and *de novo* metagenomic assembly. Viral identification and dereplication at 95% average nucleotide identity (ANI) over 85% alignment fraction (AF) resulted in a library of 49,745 viral operational taxonomic units (vOTUs) ≥3kb in length. Functional potential of a non-redundant catalog of 485,806 protein clusters (95% average amino acid identity [AAI], 90% AF) was evaluated through annotation against the KOfam database.

This integrated dataset represents one of the most comprehensive descriptions of the healthy infant gut virome to date, providing the statistical power necessary to resolve key developmental and functional milestones which may be obscured in smaller, single-cohort studies.

### The infant gut virome is highly individualized but with conserved bacterial hosts

To assess the distribution of vOTUs across infant gut viromes, we calculated their prevalence in five discrete age bins based on sampling density and their alignment with key infant developmental milestones: 0 months (neonatal), 1-3 months (early infancy), 4-6 months (transitional), 7-12 months (late infancy), >1 year (toddler). In general, vOTU prevalence, defined as the fraction of infants carrying a given vOTU, was low, with a long tail of rare vOTUs (**Figure 2A**). The maximum prevalence for any single vOTU remained low across all developmental stages, ranging from 15.4% (at 1-3 months) to 48% (at 4-6 months). Notably, no single vOTU was ever prevalent in more than half of the infants in any age bin, suggesting a lack of a core virome signature at the sequence-level.

**Figure 2.**
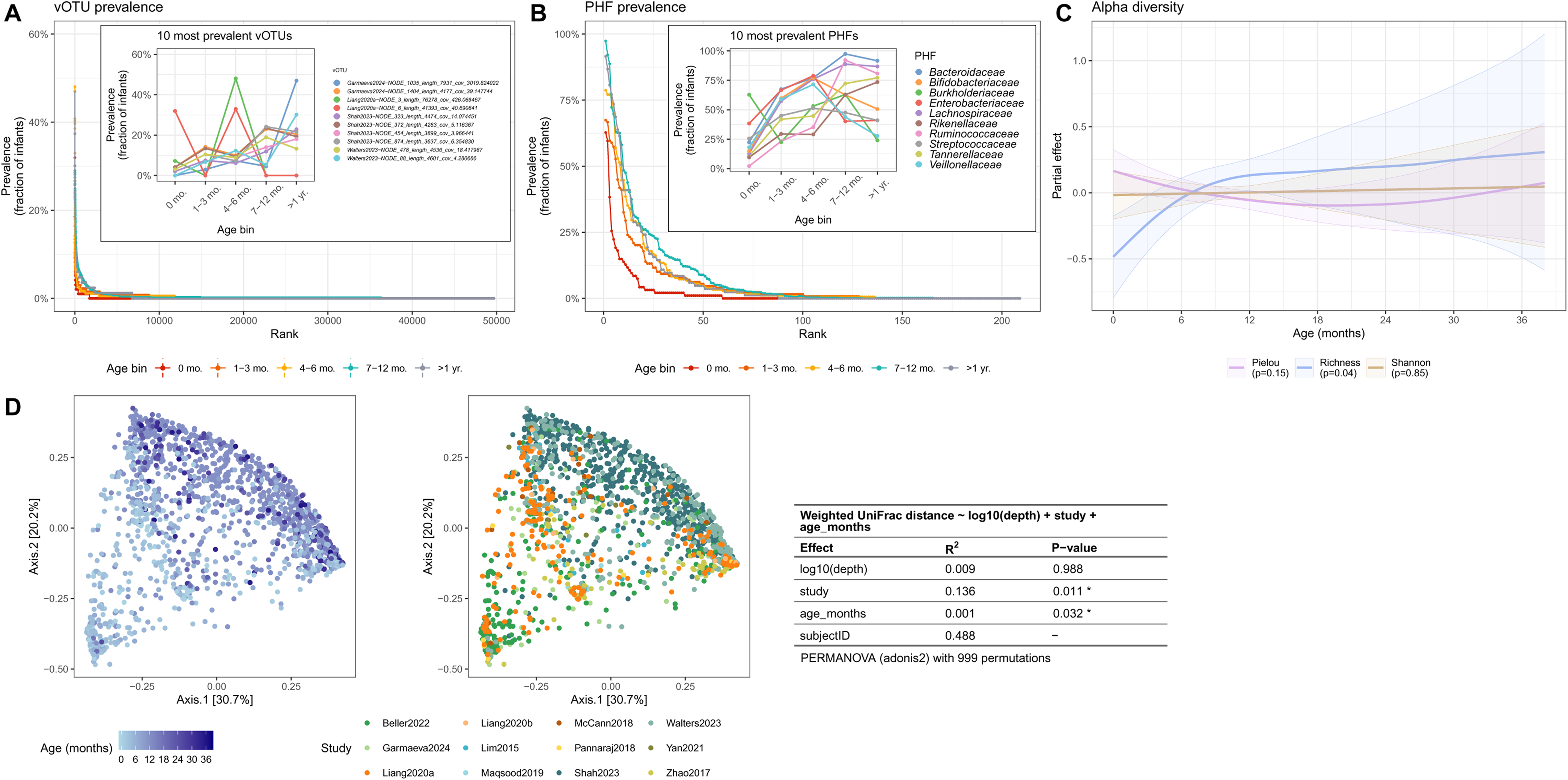
Prevalence and diversity patterns of the gut virome across the first three years of life. Prevalence of **(A)** 49,745 viral operational taxonomic units (vOTUs), and **(B)** phage host families (PHFs) across five developmental age bins. Inset plots highlight the mean prevalence of the top ten most prevalent vOTUs and PHFs. **(C)** Longitudinal alpha diversity trajectories modelled using generalized additive mixed models (GAMMs). The fitted smooth represents the non-linear relationship between each metric and infant age (in months), where the shaded ribbon represents the 95% confidence interval. **(D)** Community structure visualized with principal coordinates analysis (PCoA) of weighted UniFrac distances at the PHF level. Samples are coloured according to infant age (in months, left) and original study cohort (right). Accompanying PERMANOVA results quantify the relative contribution of metadata variables to overall virome beta diversity.

While this highlights the individuality of the infant virome, we hypothesized that more consistent patterns may emerge at a broader level, where unique vOTUs may converge upon a core set of shared bacterial hosts. Phage host families (PHFs), groupings of vOTUs based on their predicted bacterial host family, provide a functionally relevant unit of classification that can improve such comparisons (20). After grouping the data at the PHF level, prevalence increased substantially (**Figure 2B**). The top PHFs in each age bin were found in a majority of infants, reaching as high as 97.5% for *Bacteroidaceae* at 7-12 months. We also observed clear temporal successions in host targets. For example, phages targeting *Bifidobacteriaceae*, a dominant bacterial family in the neonate gut and key consumer of human milk oligosaccharides (13), reached peak prevalence during the first 6 months of life, mirroring the typical period of exclusive milk consumption. Across all age bins, the number of core PHFs detected in more than half of infants, increased steadily from 2 at birth to 9 in toddlers (>12 months).

Together, these results highlight the importance of host-based analysis, suggesting that while prevalence of individual vOTUs is highly variable, potentially limiting downstream analyses, the virome maintains functional redundancy by consistently targeting specific bacterial families throughout development.

### Gut viral diversity expands rapidly during the first year of life before stabilizing

To model vOTU-level alpha diversity trajectories while controlling for inter-study heterogeneity and repeated measures, we implemented generalized additive mixed models (GAMMs). After accounting for fixed effects (sequencing depth, study) and random effects (individual), only observed richness had a significant association with age over the entire sampling period (p = 0.042), increasing significantly during the first 8 months of life, before reaching a plateau (**Figure 2C, Figure S1**). In contrast, the Shannon index remained stable throughout the three-year sampling period. This combination of increasing richness and stable Shannon diversity implies a shift in community structure. Indeed, Pielou’s evenness decreased, though not significantly, during the first 18 months before returning to baseline levels. To evaluate between-sample diversity, we computed PHF-level beta diversity using weighted UniFrac distances (**Figure 2D**). Infant age and study of origin were both significant drivers of community composition.

### The infant gut virome converges to a common functional state

These age-related shifts in diversity prompted a deeper analysis of the dynamics of gut virome assembly. Specifically, we sought to quantify the rate of change or developmental velocity, and the level of inter-individual variation or beta dispersion to characterize how the virome stabilizes during the first years of life. GAMMs were implemented to account for fixed (sequencing depth, study) and random effects (individual).

Virome developmental velocity (VDV) was defined as the weighted UniFrac distance between two consecutive samples from the same infant, divided by the time interval (in months) between samples. VDV had a significant negative association with age (p < 0.001), highest at birth and stabilizing by approximately 6 months (**Figure 3A**). Combined with the richness findings, these data suggest that the first 6 to 8 months of life represent the most dynamic period of gut virome development.

**Figure 3.**
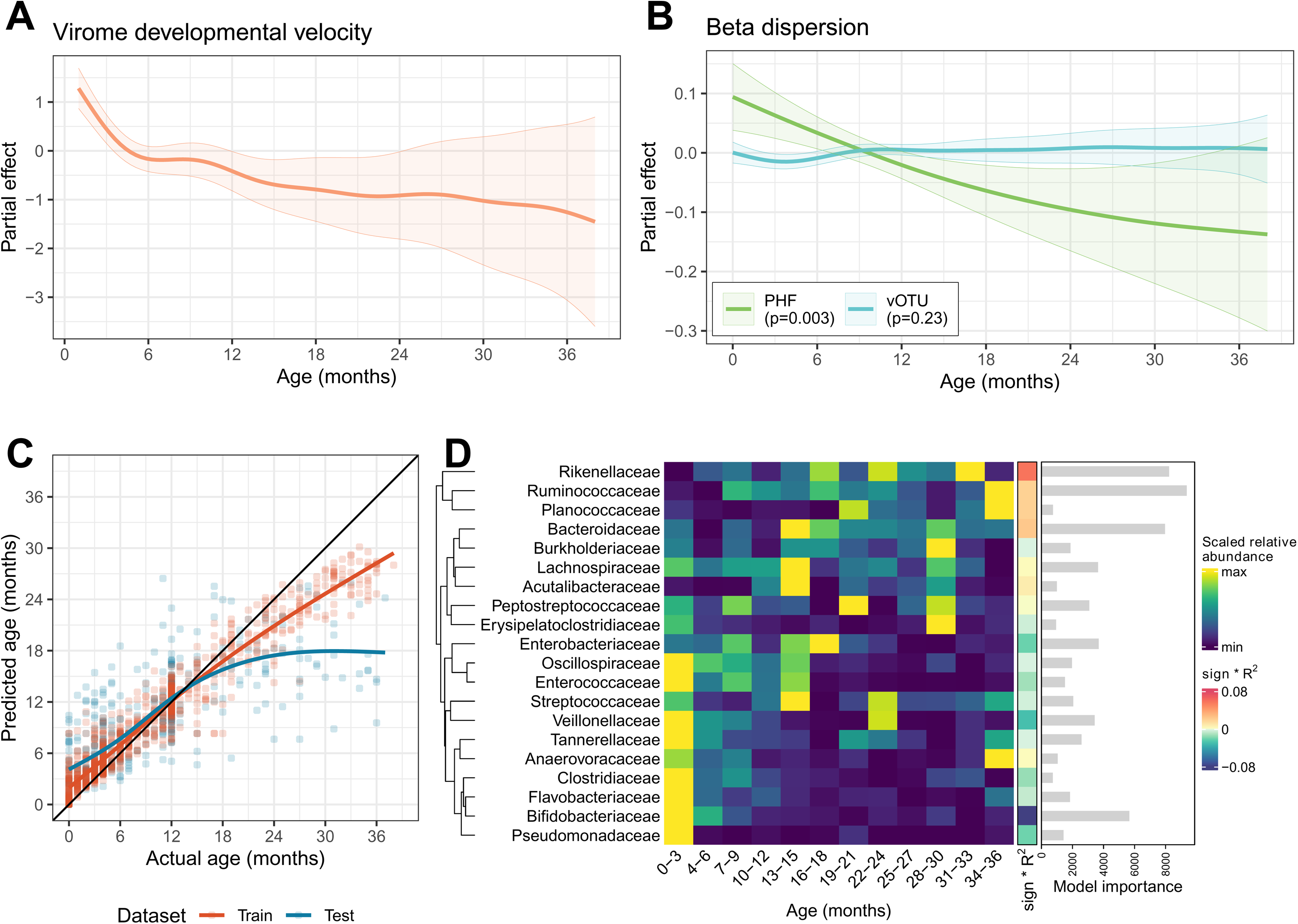
Rapid and predictable assembly of the infant gut virome. **(A)** Virome developmental velocity (VDV) modelled using a GAMM. The fitted smooth represents the non-linear relationship between VDV and infant age in months (p < 0.001), where the shaded ribbon represents the 95% confidence interval. **(B)** Beta dispersion, calculated at the vOTU and PHF levels, modelled using GAMMs. The fitted smooth represents the non-linear relationship between each model and infant age (in months), where the shaded ribbon represents the 95% confidence interval. **(C)** Random forest regression model performance for predicting infant chronological age from PHF-level virome composition. Predicted versus actual age (in months) is shown for both training and testing datasets. **(D)** Heatmap of the top twenty most important PHF features in the age predictor model, with mean relative abundance across 3-month age bins (min/max scaled for each PHF). Accompanying coloured strip and bar plot indicate relative overall contribution and importance of each PHF to the model, respectively.

To evaluate gut virome convergence and changes in inter-individual variation over time, beta dispersion was calculated as the distance of each sample to its age-specific centroid in multivariate ordination space. At the PHF level, beta dispersion significantly decreased with age (p = 0.0032), continuing this downward trend beyond 36 months (**Figure 3B**). In contrast, beta dispersion at the vOTU level had no association with age (p = 0.23), suggesting that the gut virome remains highly individualized at the sequence level throughout early life. The discordance between PHF- and vOTU-level beta dispersion patterns suggests that infant gut viromes converge toward a common structure of phage-host interactions, while remaining highly individualized at the sequence level.

### A subset of phage host families enables accurate prediction of infant age

Given the successional patterns observed thus far, we next investigated whether gut virome composition could serve as a predictor for infant age. We trained a random forest regression model to predict infant age based on PHF abundances, allowing us to identify a set of age-discriminatory features that define healthy virome maturation. A 70%/30% ratio was used to split the dataset into train and test subsets, respectively.

Performance was evaluated using root mean square error (RMSE) and R^2^ metrics. On training data, the model had high accuracy (RMSE = 2.49 months, and R^2^ = 0.93), while maintaining predictive power on the test set (RMSE = 5.14 months, R^2^ = 0.555; **Figure 3C**). Using a three-month sliding window to assess local performance, we found that model RMSE was lowest during the first 20 months of life, the period of highest sampling density and most rapid virome turnover, before increasing significantly as the virome stabilized in older infants (**Figure S2A**). Leave-one-study-out cross-validation (LOSOCV) was also performed to ensure that the identified patterns were not driven by study-specific biases. In general, removal of any single study from the training dataset reduced the resulting model’s performance, indicating that each study provides important information for capturing global successional trends in virome development (**Figure S2B**, **Figure S2C**).

To identify the drivers of the age predictor model, we ranked PHFs by their predictive importance and evaluated the direction of their associations with age (**Figure 3D**). The most age-discriminatory features included *Ruminococcaceae*, *Rikenellaceae*, *Bacteroidaceae*, and *Bifidobacteriaceae*. Early-life age prediction was primarily driven by the decline of *Bifidobacteriaceae* and *Veillonellaceae* (highest during first 3 months), while transition to toddler age was marked by increasing abundances of *Rikenellaceae* and *Bacteroidaceae* (highest after 13 months) (**Figure 3D**).

The ability to accurately predict infant age based on PHF-level community composition confirms that the gut virome follows a predictable successional pattern, likely driven by the stabilization of core bacterial hosts during this same period.

### Temperate phages dominate the extracellular gut virome during early infancy

Several studies have hypothesized that the gut virome is quickly populated after birth by the induction of temperate phages from colonizing bacteria, though the specific triggers remain unknown. To test this hypothesis, we predicted vOTU lifestyles and modelled the cumulative relative abundance of temperate phages across the first few years of life, using GAMMs to account for fixed (sequencing depth, study) and random effects (individual). Temperate phage abundances were significantly associated with age (p = 0.002), highest at birth and decreasing rapidly thereafter (**Figure 4A**).

**Figure 4.**
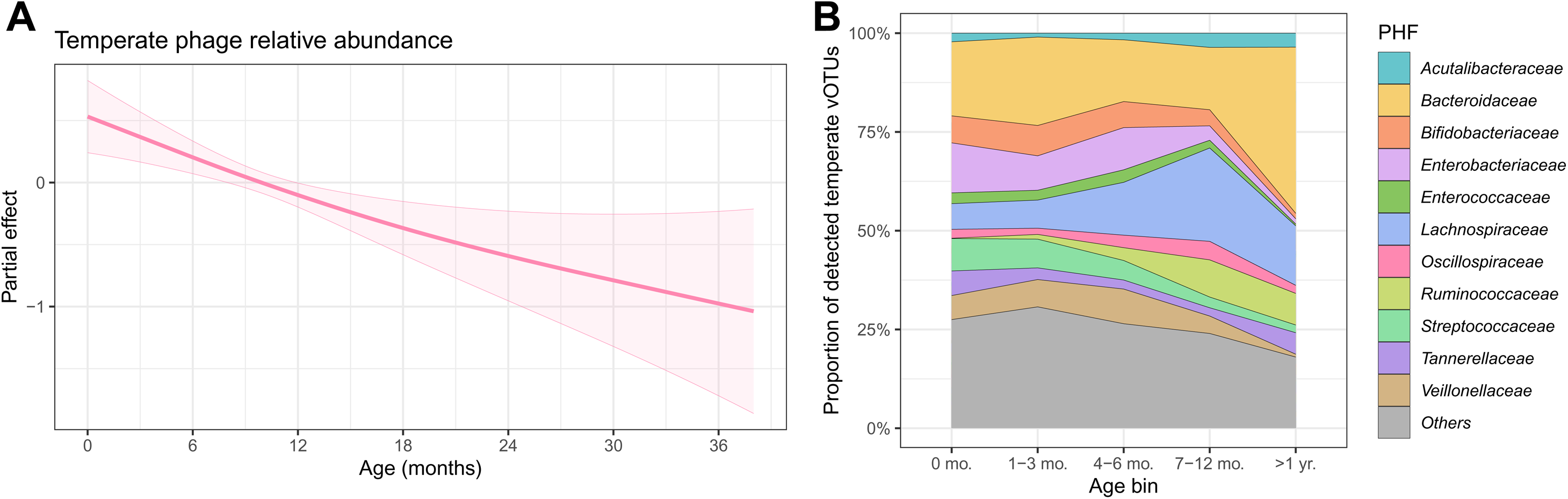
Abundance and diversity of the temperate infant gut virome. **(A)** Cumulative temperate phage relative abundance modelled using a GAMM. The fitted smooth represents the non-linear relationship between temperate phage relative abundance and infant age in months (p = 0.002), where the shaded ribbon represents the 95% confidence interval. **(B)** Composition of the temperate virome at the PHF level across five developmental age bins, representing the compositional richness (proportion of unique vOTUs) within each bin, unweighted by read abundance.

The compositional diversity of the temperate virome also varied with time. At birth, most of the detected temperate vOTUs were predicted to infect *Bacteroidaceae*, *Enterobacteriaceae*, *Bifidobacteriaceae*, and *Streptococcaceae* hosts (**Figure 4B**). By one year of age, the temperate virome was primarily composed of phages infecting *Lachnospiraceae*, *Bacteroidaceae*, and *Ruminococcaceae*, reflecting changes in bacteriome composition over this same period.

### Functional specialization of the gut virome mirrors key developmental milestones

To further characterize the functional potential of the infant virome, protein-coding sequences were predicted from the vOTU library and then clustered into a non-redundant catalog based on ≥95% average amino acid identity (AAI) and ≥90% coverage (**Figure 1C**). A total of 797,941 protein-coding sequences were predicted from the library of 49,745 vOTUs, resulting in 485,806 non-redundant protein clusters (reduction of 39.1%). We performed functional annotation of these non-redundant gene catalog against the KOfam database and filtered for auxiliary metabolic genes (AMGs) using a publicly available curated list of KEGG orthologs (KOs) (54).

We conducted evaluated enrichment of KEGG pathway categories among PHFs, to determine if a functional category is significantly overrepresented relative to the size of that PHF. Among the 10 most prevalent PHFs, *Ruminococcaceae* and *Bacteroidaceae* PHFs were the most functionally diverse PHFs, enriched in 37 and 30 pathway categories, respectively (**Figure 5**). These AMGs included genes for carbohydrate metabolism, biosynthesis of secondary metabolites, and amino acid metabolism.

**Figure 5.**
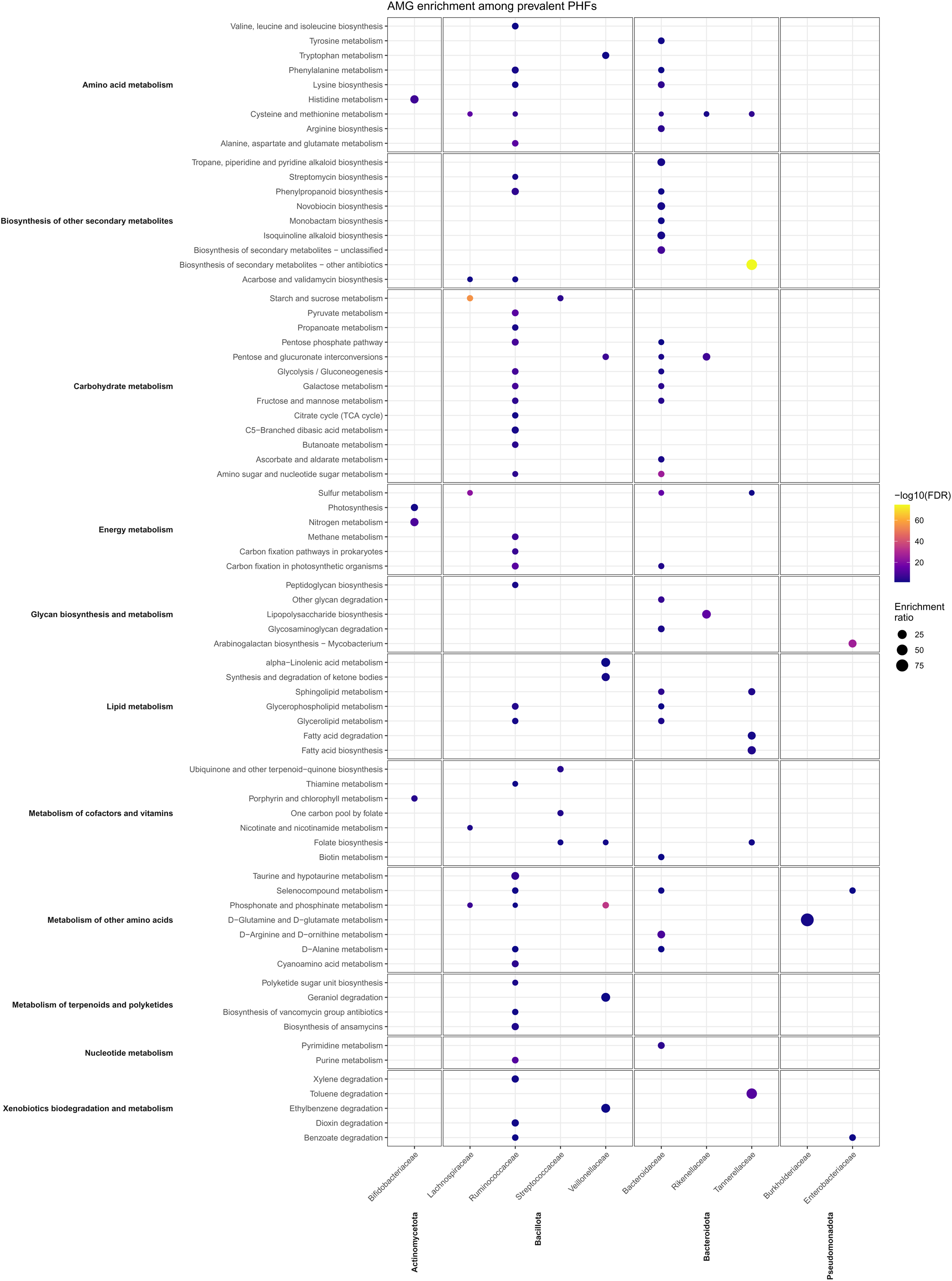
Functional specialization of the infant gut virome through auxiliary metabolic gene enrichment. Enrichment of KEGG pathway categories among the 10 most prevalent PHFs. KEGG pathways (y-axis) are organized by broader KEGG metabolism categories. Dot size indicates the enrichment ratio, while colour indicates statistical significance. Enrichment was determined using a hypergeometric test (Fisher’s exact test) to determine identify significantly overrepresented functional categories relative to the total gene count and size of a PHF. Only significant (FDR < 0.05) results are shown.

The relative abundance of different AMGs also followed a temporal succession, reflecting the evolving requirements of the infant gut (**Figure S3**). The neonatal period (0-3 months) was characterized by a high abundance of genes involved in energy, glycan, and amino acid metabolism. As infants transitioned to solid food (7-12 months), we observed a peak in AMGs associated with carbohydrate, lipid, xenobiotic, and nucleotide metabolism, which is sustained into the third year of life. We attempted to classify vOTUs into PHFs using a random forest classification model based on AMG profiles (presence/absence) and age at detection. The model’s poor performance (mean F1 score across all classes/PHFs of 0.58) suggests a high degree of functional redundancy across PHFs, where specific metabolic repertoires are not necessarily informative of host.

## Discussion

In recent years, there has been growing interest in the human gut virome, especially in the context of early life development. Here, we conducted a meta-analysis of twelve infant cohorts across 8 countries, with 1,893 samples from 999 infants, to model gut virome trajectories during the first three years of life. Although the studies have each individually identified patterns within their respective datasets, geographical and analytical differences have hindered cross-study comparisons. Using a unified bioinformatic pipeline and statistical modelling to control for cohort-and individual-specific effects, we identified significant increases in viral richness during the first eight months of life. By grouping phages together by the hosts family (PHF) they are predicted to infect, we could overcome the issue of virome interindividuality to resolve common successional trends (20). We observed high prevalence of multiple PHFs among infants, with *Bacteroidaceae* phages being the most prevalent (detected in 97.5% of infants at 7-12 months). Conversely, the most prevalent vOTU was detected in only 48% of infants at a single timepoint (4-6 months). This contrast suggests that while vOTUs are highly individualized, their ecological niches (at a phage-host family level) are conserved across individuals and cohorts.

To evaluate virome temporal dynamics, we define a novel metric, the virome developmental velocity (VDV). VDV was highest after birth and decreased with time, stabilizing by six months of age. Similarly, beta dispersion, a measure of sample relatedness, decreased with time, suggesting that infant gut viromes converge toward a common structure, despite remaining highly individualized at the sequence level. Taken together with the richness results, these data suggest that the first year of life represents the most active phase of gut virome development, with high viral turnover and continuous introduction of new taxa, likely mirroring similar major shifts in gut bacteriome composition during this same period.

Using a machine learning approach, we identified several predictors of healthy virome development. Phages targeting *Ruminococcaceae*, *Rikenellaceae*, *Bacteroidaceae*, and *Bifidobacteriaceae* were among the most important predictors of infant age. Alongside these changes in virome structure is a clear shift in phage-host interactions as the gut environment stabilizes.

Consistent with other studies, we observed decreasing abundances of temperate phages with time, suggesting a transition to an adult-like microbiota where temperate phages primarily exist as prophages within host genomes (55). While a decrease in temperate phage abundance implies a corresponding increase in the lytic fraction, we adopted a conservative analytical approach by focusing exclusively on high-confidence temperate vOTUs. Indeed, a temperate phage missing key lysogeny genes due to incomplete metagenomic assembly may falsely be classified by computational tools as being lytic.

Beyond changes in phage lifestyle, the functional potential of the gut virome also experiences significant shifts in early life. The high enrichment of AMGs within phages targeting *Ruminococcaceae* and *Bacteroidaceae* – two bacterial families essential for fermentation of dietary fibers (13) – suggests that the extracellular virome provides a metabolic toolkit which supports host adaptation to a post-weaning, plant polysaccharide-rich diet, potentially enhancing their competitive fitness during dietary transitions.

It is important to note that all twelve studies are from the Global North, with many from high-income countries. Further virome studies from more diverse geographic regions and low- and middle-income countries are necessary to gain a better understanding of global patterns in virome development. Furthermore, all studies included here use library preparation methods which are heavily biased towards sequencing of double-stranded DNA, precluding insights into RNA and single-stranded DNA viruses. Inconsistent metadata collection and reporting across studies also limited the analyses we could perform. For example, only 21% of samples had associated sex and delivery mode information, significantly limiting the statistical power of our models if we were to train using only this small subset.

Ultimately, by integrating twelve global cohorts, our work characterizes healthy gut virome development in early life, highlighting a conserved successional program and baseline for future studies on the viral drivers of human health.

## Acknowledgements

This work was funded by a Canadian Institutes of Health Research Canada Project grant (PJT-175065), Natural Sciences and Engineering Research Council of Canada Discovery Grant (RGPIN-2023-04216), and Canada Research Chair in Gut Microbial Interactions (Tier 2) to Corinne F. Maurice. Michael Shamash is supported by the Canadian Institutes of Health Research Canada Graduate Scholarship to Honor Nelson Mandela (CIHR CGS-D; #DF2-187718), and the *Fonds de recherche du Québec-Santé: Bourse de formation au doctorat* (FRQS; #311071).

## Data and code availability

Code used for data analysis is available at https://github.com/mshamash/infant_gut_virome_manuscript.

## Author contributions

M.S. curated data for analysis, conceived and performed the analyses, prepared figures and tables, authored and reviewed drafts of the manuscript, and approved the final draft. C.F.M. conceived the analyses, reviewed drafts of the manuscript, and approved the final draft.

## Declaration of generative AI and AI-assisted technologies in the writing process

During the preparation of this work the authors used Google Gemini (Alphabet Inc., Mountain View, USA) to improve sentence clarity and readability. Overall sentence meaning was not altered, and there were no changes to any of the presented data, facts, or conclusions by using this tool. After using this tool, the authors reviewed and edited the content as needed and take full responsibility for the content of the published article.

## Supplementary Table and Figure Legends

**Supplementary Table S1. Summary of metadata and sampling information for the 12 integrated cohorts.** Breakdown of the studies included in this meta-analysis, including sampling location, cohort size (number of participants & number of samples), and the age range of sampling. The study design for each study is abbreviated as follows: L (longitudinal) or C (cross-sectional).

**Supplementary Figure S1. Rate of change of gut viral richness during early life.** The first derivative of the generalized additive mixed model (GAMM) for observed richness (see **Figure 2C**) was calculated with respect to infant age in months. The shaded ribbon represents the 95% confidence interval. The vertical dashed line at x = 7.68 months indicates the earliest point at which the confidence interval crosses y = 0. This threshold represents the point at which the rate of richness accumulation is no longer significantly different from zero, indicating that virome richness has plateaued.

**Supplementary Figure S2. Validation of the age predictor model. (A)** Model performance throughout development, evaluated as the root mean square error (RMSE) within 3-month sliding windows. Points indicate the RMSE for each window, while bars indicate the number of sample size (n) within that 3-month period. **(B)** Comparison of model accuracy across the 12 cohorts evaluated via leave-one-study-out cross-validation (LOSOCV). Red bars (full model) represent the RMSE when the model trained on the entire original training dataset is tested on a specific cohort. Blue bars (LOSOCV) represent the RMSE when the model is trained excluding that specific cohort and then tested on it. **(C)** Age predictions from both the full regression model and LOSOCV permutations.

**Supplementary Figure S3. Functional maturation of the infant gut virome.** Heatmap of the mean relative abundance of KEGG metabolism categories across 3-month age bins. To highlight successional shifts, values are min/max scaled across each category (row).

