## Supplementary figures and images for "The infant gut virome assembles rapidly and predictably over the first three years of life"

### Supplemental Figure 1

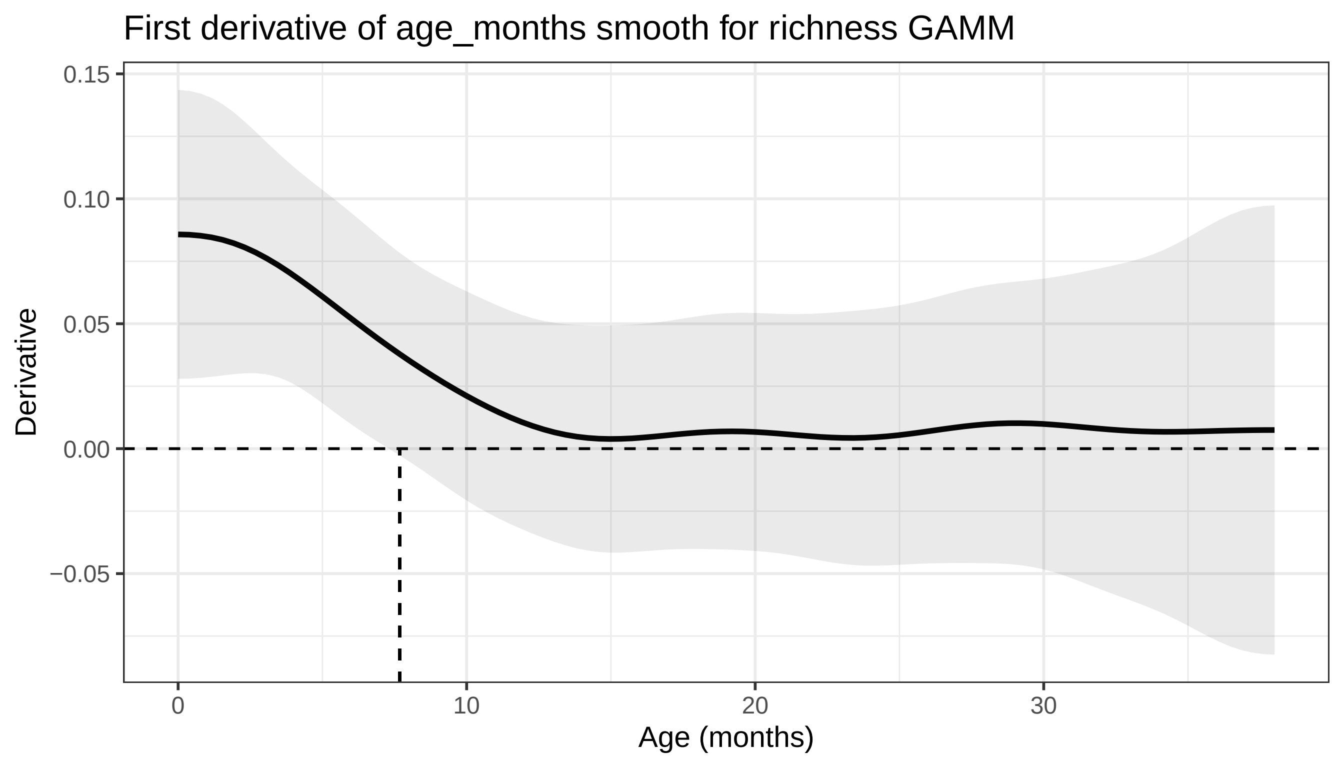

### Supplemental Figure 2

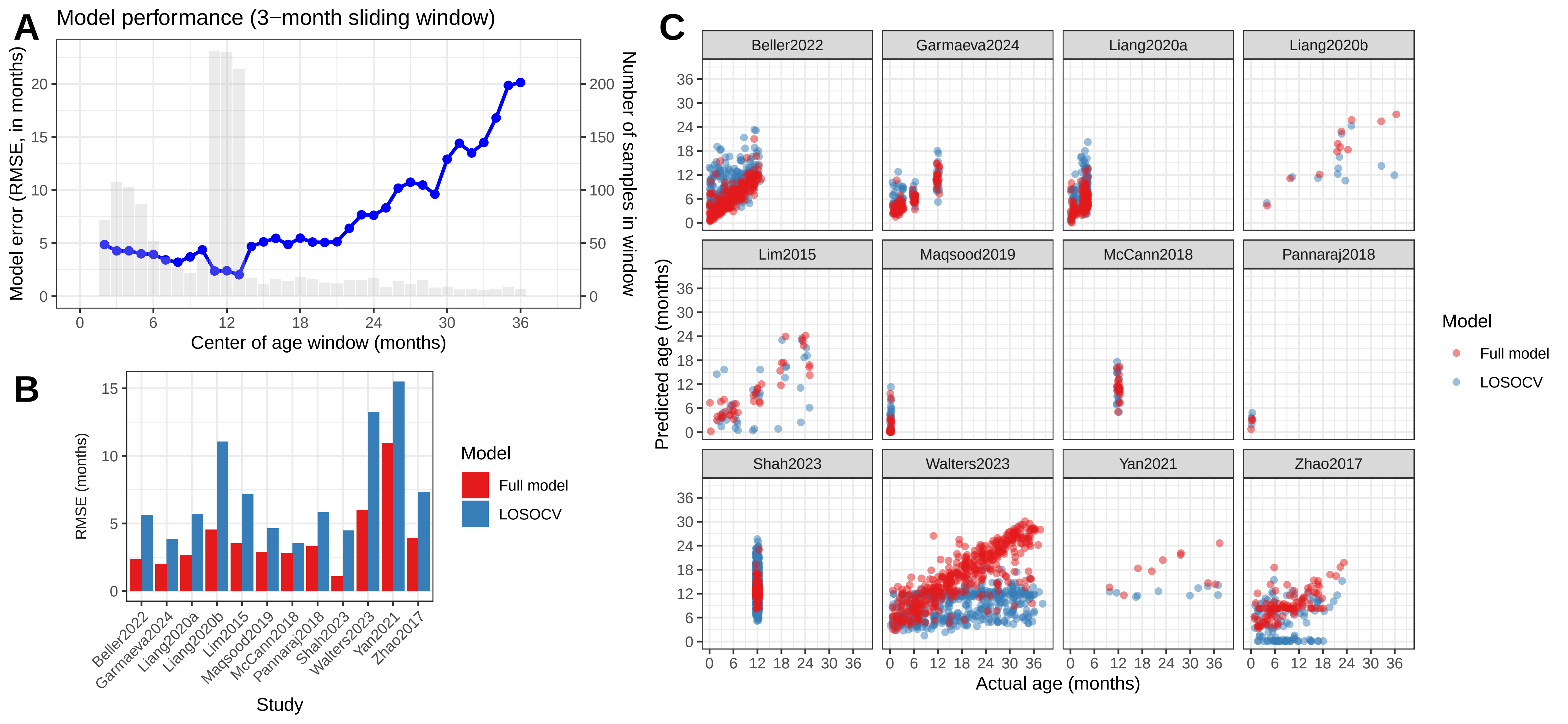

### Supplemental Figure 3

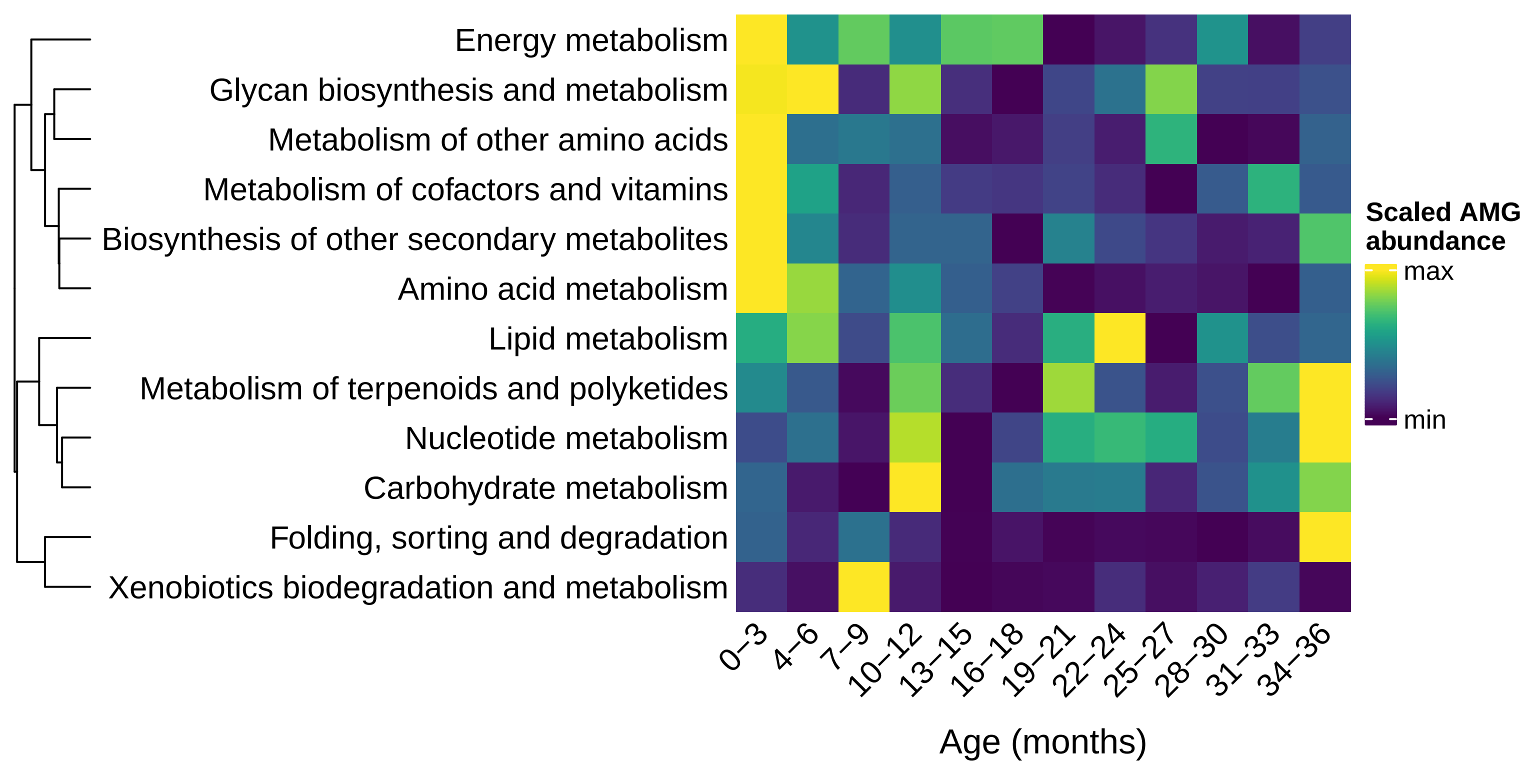
