## Supplemental Table 1 for "The infant gut virome assembles rapidly and predictably over the first three years of life"

| Study | Ref. | Accession | Sampling location(s) | Sampling period (months) | Study design | Number of participants | Number of samples |
|---|---|---|---|---|---|---|---|
| Lim2015 | 23 | PRJNA284162 | United States | 0 - 24 | L | 8 | 48 |
| Zhao2017 | 24 | PRJNA387903 | Finland, Estonia | 1 - 23 | L | 11 | 106 |
| McCann2018 | 25 | PRJNA385126 | Ireland | 12 | C | 20 | 20 |
| Maqsood2019 | 26 | PRJEB33578 | United States | 0 | C | 51 | 51 |
| Liang2020a | 11 | PRJNA524703 | United States | 0 - 4 | L | 143 | 184 |
| Beller2022 | 28 | PRJNA693793 | Belgium | 0 - 13 | L | 8 | 304 |
| Walters2023 | 29 | PRJNA916952 | United States | 0 - 38 | L | 51 | 421 |
| Shah2023 | 30 | PRJEB46943 | Denmark | 12 | C | 647 | 647 |
| Garmaeva2024 | 22 | Not publicly available | Netherlands | 1 - 12 | L | 29 | 81 |
| Pannaraj2018 | 31 | PRJNA427187 | United States | 0 | C | 10 | 10 |
| Yan2021 | 32 | PRJNA641593 | China | 12 - 36 | C | 10 | 10 |
| Liang2020b | 27 | PRJNA524703 | United States | 4 - 36 | C | 11 | 11 |
| | | | | | Total | 999 | 1,893 |
